# Large-scale phenome-wide association study of *PCSK9* loss-of-function variants demonstrates protection against ischemic stroke

**DOI:** 10.1101/210302

**Authors:** Abhiram S. Rao, Daniel Lindholm, Manuel A. Rivas, Joshua W. Knowles, Stephen B. Montgomery, Erik Ingelsson

**Affiliations:** Department of Bioengineering, Stanford University, Stanford, CA 94305, USA; Department of Medical Sciences, Cardiology, Uppsala University, Uppsala, Sweden (DL); Uppsala Clinical Research Center, Uppsala, Sweden; Department of Biomedical Data Science, Stanford University, Stanford, CA 94305, USA; Division of Cardiology, Department of Medicine, Stanford University, Stanford, CA 94305, USA; Department of Genetics, Stanford University School of Medicine, Stanford, CA 94305, USA; Department of Pathology, Stanford University School of Medicine, Stanford, CA 94305, USA; Division of Cardiovascular Medicine, Department of Medicine, Stanford University, Stanford, CA 94305, USA

## Abstract

*PCSK9* inhibitors are a potent new therapy for hypercholesterolemia and have been shown to decrease risk of coronary heart disease. Although short-term clinical trial results have not demonstrated major adverse effects, long-term data will not be available for some time. Genetic studies in large well-phenotyped biobanks offer a unique opportunity to predict drug effects and provide context for the evaluation of future clinical trial outcomes. We tested association of the *PCSK9* loss-of-function variant rsll591147 (R46L) in a hypothesis-driven 11 phenotype set and a hypothesis-generating 278 phenotype set in 337,536 individuals of British ancestry in the United Kingdom Biobank (UKB), with independent discovery (n = 225K) and replication (n = 112K). In addition to the known association with lipid levels (OR 0.63) and coronary heart disease (OR 0.73), the T allele of rs11591147 showed a protective effect on ischemic stroke (OR 0.61, p = 0.002) but not hemorrhagic stroke in the hypothesis-driven screen. We did not observe an association with type 2 diabetes, cataracts, heart failure, atrial fibrillation, and cognitive dysfunction. In the phenome-wide screen, the variant was associated with a reduction in metabolic disorders, ischemic heart disease, coronary artery bypass graft operations, percutaneous coronary interventions and history of angina. A single variant analysis of UKB data using TreeWAS, a Bayesian analysis framework to study genetic associations leveraging phenotype correlations, also showed evidence of association with cerebral infarction and vascular occlusion. This result represents the first genetic evidence in a large cohort for the protective effect of *PCSK9* inhibition on ischemic stroke, and corroborates exploratory evidence from clinical trials. *PCSK9* inhibition was not associated with variables other than those related to low density lipoprotein cholesterol and atherosclerosis, suggesting that other effects are either small or absent.

## Background

Cardiovascular disease (CVD) accounts for over a quarter of annual deaths in the United States, and is the most common cause of mortality globally.^1,2^ Hypercholesterolemia has long been established as one of the most important risk factors for the development of CVD;^3^ statin-based therapeutic regimens reduce low-density lipoprotein cholesterol (LDL-C), thereby decreasing CVD and all-cause mortality.^4–6^ Statins achieve their LDL-C lowering effects by inhibiting 3-hydroxy-3-methylglutaryl-coenzyme A reductase (*HMGCR*)^7^ and have become amongst the most widely prescribed drugs across all drug classes. However, many patients on medium-to high-intensity statin therapy are unable to achieve a clinically recommended reduction in LDL-C, leading to the consideration of other LDL-C lowering therapies.^8,9^

*PCSK9* (proprotein convertase subtilisin/kexin type 9) inhibitors are an important new class of LDL-C lowering drugs that were discovered based on human genetic studies.^10,11^ The gene product of *PCSK9* binds to the low-density lipoprotein receptor (*LDLR*) and promotes its degradation, thereby increasing levels of circulating LDL-C.^12^ Genetic studies have established the association of variants in the *PCSK9* locus with hypercholesterolemia and atherosclerotic CVD (ASCVD),^10,13,14^ and fine-mapping efforts have revealed multiple independent signals within the locus.^15,16^ *PCSK9* variants can independently influence lipid levels^11,17^ and also act to modify therapeutic response to statins.^18,19^ Early evidence of this association generated interest in *PCSK9* as a potential drug target,^20^ and two recent large-scale clinical trials have confirmed the protective effects of the monoclonal antibody *PCSK9* inhibitors evolocumab and alirocumab on composite CVD and hypercholesterolemia endpoints.^21,22^ *PCSK9* inhibitors are currently indicated for patients with ASCVD or Familial Hypercholesterolemia (FH) who are unable to achieve optimal LDL-C on statin-based regimens,^23^ but questions remain about the potential adverse effects and cost-effectiveness of these drugs. While existing clinical studies show that they are generally well-tolerated in the short term,^21,24^ there remain questions about the long term effects of *PCSK9* inhibitors on type 2 diabetes (T2D) and cognitive dysfunction, among other phenotypes.^25–27^

A phenome-wide association study (PheWAS) of *PCSK9* loss-of-function (LoF) variants has the potential to predict adverse effects of *PCSK9* inhibition and generate hypotheses about novel indications. Comparing carriers of such LoF variants with non-carriers can simulate the effects of blocking the corresponding protein. Hence, this approach can offer insights into beneficial as well as adverse drug effects that are likely to result from direct inhibition of *PCSK9,* the intended drug target; predicting antibody-related adverse effects is more difficult. In the present study, we combined a hypothesis-driven analysis of 11 phenotypes defined *a priori* based on concurrent literature with a PheWAS of *PCSK9* LoF variants in 337,536 individuals of British ancestry from the United Kingdom Biobank (UKB). We leveraged the increase in power offered by this focused hypothesis-driven testing and applied newly-developed methods that use the correlation structure of phenotypes to reduce the multiple testing burden in our phenome-wide screen.

## Methods

### Study sample

The UKB is a prospective cohort study with genotype and detailed phenotype information for up to 502,628 individuals aged 40-69 years when recruited between 2006 and 2010. The phenotype data includes hospital diagnoses, self-reported data based on questionnaires, interview-based data, and physical and functional measurements. The genotype data includes positions that are either directly genotyped on one of two arrays (UKB Axiom and UK BiLEVE arrays), or imputed using a reference panel from the Haplotype Reference Consortium (HRC). The study was approved by the North West Multi-Centre Research Ethics Committee and all participants provided written informed consent to participate. For the present study, we used a study sample of 337,536 unrelated individuals of British ancestry after excluding six individuals who chose to withdraw their data from the UKB.

### Phenotypes studied

We compiled two sets of phenotypes to study associations between *PCSK9* loss-of-function variants and diverse outcomes across the whole phenome. First, we created a hypothesis-driven set of 11 phenotypes informed by concurrent literature,^12,17,21,26,27^ including cholesterol medication status, coronary heart disease, all stroke, ischemic stroke, hemorrhagic stroke, T2D, cataracts, heart failure, atrial fibrillation, epilepsy, and a cognitive function test (detailed definitions in Table S1). Next, we created a hypothesis-generating phenome-wide set comprising blocks of ICD-10 codes, groups of procedure (OPCS-4) codes, self-reported variables reflecting medical information, symptoms and environmental factors, and physical and functional measurements. We excluded variables encoding congenital malformations, injuries, and accidents, and any other binary variable definitions with fewer than 500 total cases among 337,536 individuals (Supplementary Materials). Power curves^28^ for binary phenotypes are shown in Figure S1. The final phenome-wide set included 278 phenotypes (Table S2).

### Phenome-wide association testing

Hypothesis-driven and hypothesis-generating groups of association tests with the T allele of rs11591147 (R46L, 1.7% MAF in white/European) were carried out sequentially. We randomly split the data into discovery (2/3) and replication (1/3) subsets for each phenotype, maintaining an approximately equal proportion of cases and variant carriers in both subsets for binary phenotypes. Bonferroni and Benjamini-Hochberg (FDR 5%) methods were used for multiplicity adjustment in the hypothesis-driven and hypothesis-generating sets respectively; a nominal significance threshold of 0.05 was used for replication. All tests (independent of replication) were re-analyzed in the full data to obtain more exact effect size estimates. Logistic and linear regression models were implemented using the glm() and lm() functions respectively in R (v3.3.0), and were adjusted for age, gender, ten genotype principal components and genotyping array.

### Discovering specific phenotype definitions underlying associations using Tree-WAS

Variables used in the PheWAS described above represent a combination of ICD-10, OPCS-4 or other codes to increase power and reduce the multiple-testing burden. To obtain greater detail for associations at the level of individual ICD-10 codes, we applied a recently published method called TreeWAS^29^ using a comprehensive phenome map of ICD-10 codes across 337,536 individuals that we constructed using the natural tree-like hierarchy of ICD-10 classifications. The phenome map contained 1,455 total nodes organized in three levels of hierarchy terminating with 1,262 individual ICD-10 codes. We first evaluated the evidence for association of rsll591147 across the phenome map, and similar to Cortes et al.,^29^ used a posterior probability of association (PPA) threshold of 0.75 to evaluate significance. In addition to a single variant analysis, we ran a phenome-wide screen of a genetic score created using six *PCSK9* variants (rs11591147, rs11583680, rs505151, rs562556, rs11206510, and rs2479409; Table 1; Figure S2; Supplementary Materials). The weight for the j^th^ variant in the score was drawn from a beta(1,25) distribution as *w*_*j*_ = beta(MAFj, 1,25), which is a modified version of a previously published weighting method.^30^

**Table 1:** Variants used in the weighted *PCSK9* genetic score. TreeWAS results for this score are summarized in Figure S4.

| Variant ID | Ref/Alt | Molecular impact | MAF |
| --- | --- | --- | --- |
| rs11591147 | G/T | Missense (R46L) | 0.01 |
| rs11583680 | C/T | Missense (A63V) | 0.18 |
| rs505151 | A/G | Missense (E670G) | 0.03 |
| rs562556 | A/G | Missense (V474I) | 0.17 |
| rs11206510 | T/C | Non-coding | 0.18 |
| rs2479409 | G/A | Non-coding | 0.35 |

## Results

We performed PheWAS of the *PCSK9* missense variant rs11591147 (R46L, G/T) in 337,536 individuals of British ancestry in the UKB, beginning with association tests in a 11-phenotype hypothesis-driven set, and subsequently testing phenome-wide association across 278 manually curated phenotypes. In the hypothesis-driven set, the rs11591147 T allele showed a protective effect on cholesterol medication status, coronary heart disease and ischemic stroke; Bonferroni correction was used in the discovery subset, and a nominal threshold of 0.05 was used for replication (Figure 1). We re-tested all phenotypes in the full dataset of 337,536 individuals to obtain better effect size point estimates and screen for suggestive, but non-signihcant associations; power estimates for all phenotypes are included in Table S1. We observed a nominally significant association with all stroke in the full dataset (Figure 1). In the hypothesis-generating phenome-wide analysis, the variant was significantly associated with a reduction in metabolic disorders, ischemic heart disease, coronary artery bypass graft operations, percutaneous coronary interventions, history of angina and cholesterol medication status (Figure 2). The association with history of myocardial infarction was significant in the discovery subset, but did not formally replicate using our a priori defined thresholds (see Methods). We observed nominally significant associations (p-value threshold 0.05) with 19 other variables in the hypothesis-generating set (Figure S3).

**Figure 1:**
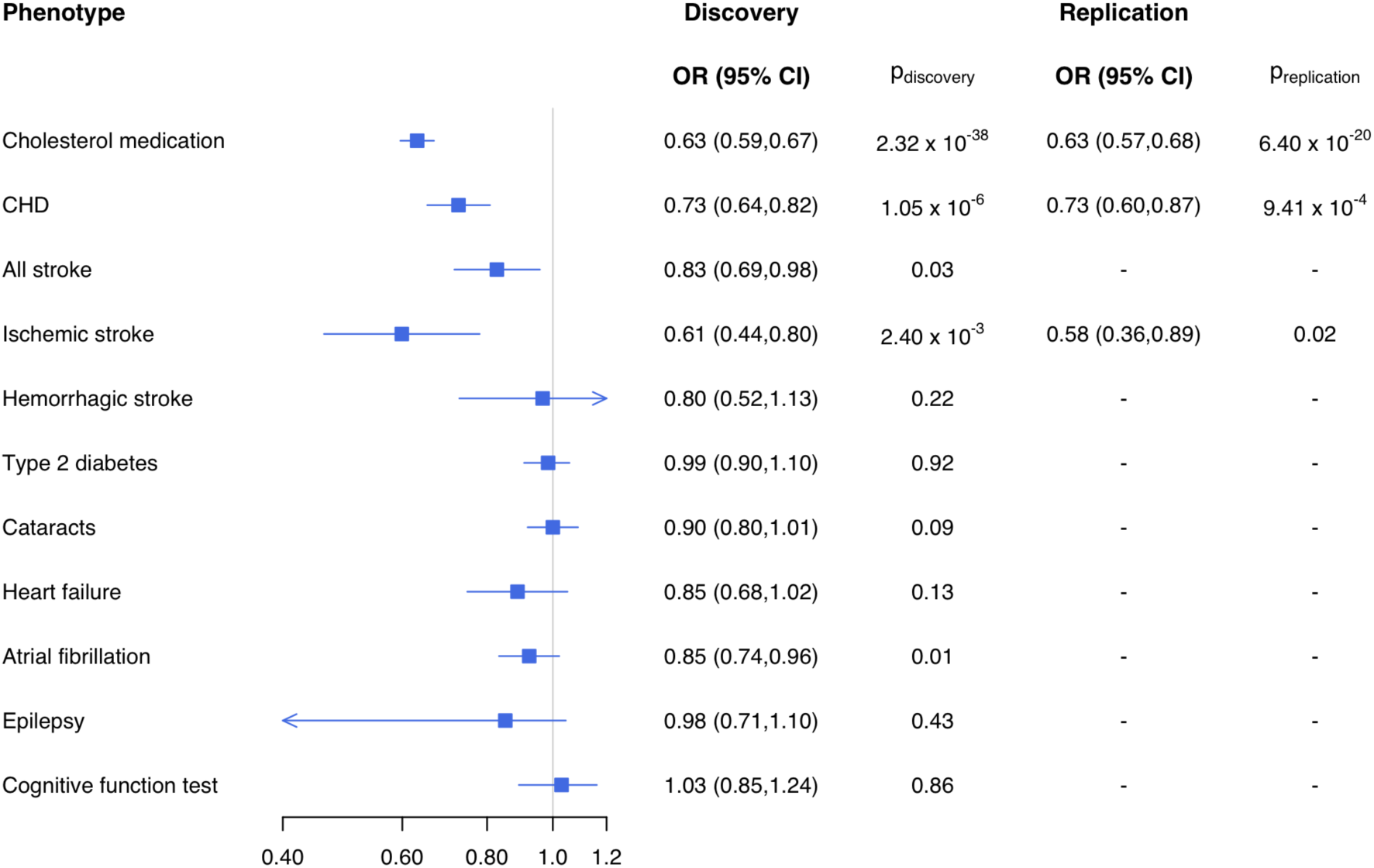
Summary of associations in the hypothesis-driven phenotype set. Forest plot estimates are based on re-analysis in the full dataset. Discovery and replication p-values are unadjusted.

**Figure 2:**
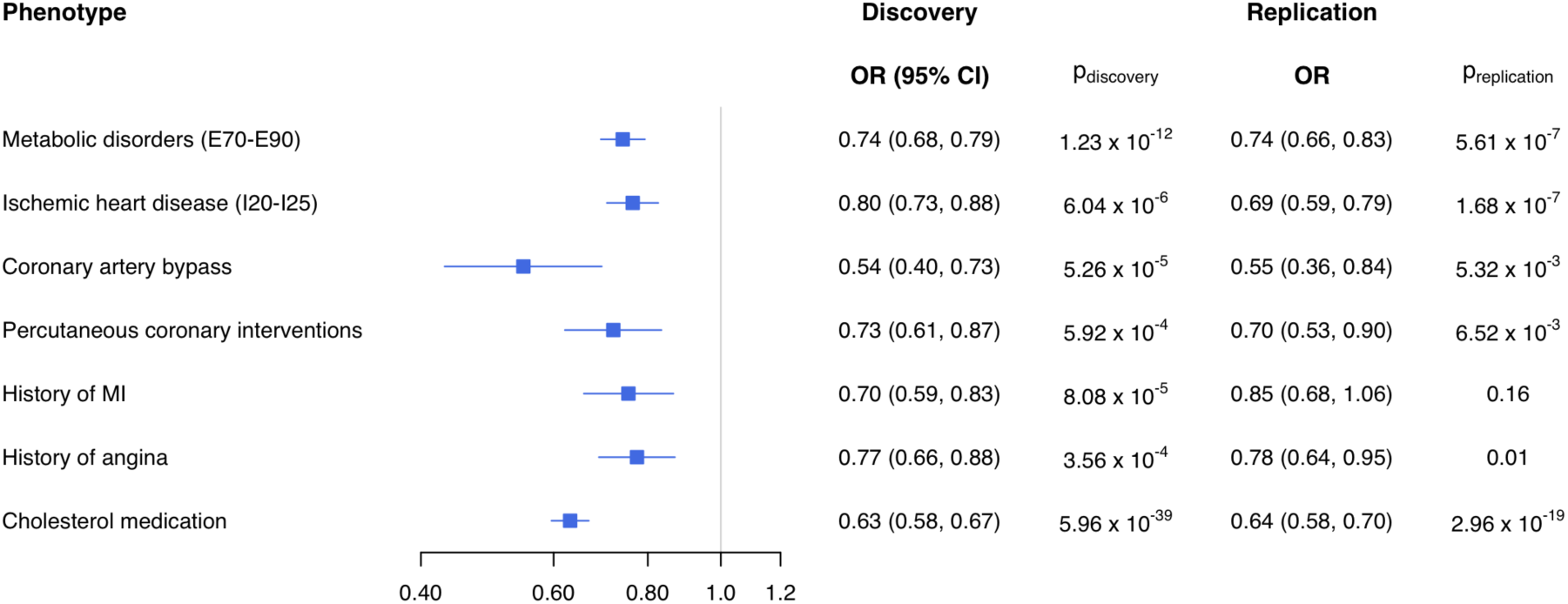
Summary of associations in the hypothesis-generating phenotype set. Forest plot estimates are based on re-analysis in the full dataset. Discovery and replication p-values are unadjusted.

We then ran a phenome-wide association screen for rs11591147 using TreeWAS with a PPA significance threshold of 0.75. We observed that the association with metabolic diseases was driven by disorders of lipoprotein metabolism (E78) and the association with ischemic heart disease was driven by all individual ICD-10 codes comprising that phenotype (I20125) except complications following myocardial infarction (123). Cerebrovascular phenotypes that showed association in the single variant analysis were primarily related to infarction (163) and occlusion (165). We did not observe significant or suggestive associations across 15 other disease categories. In total, we observed 11 active (significantly associated) nodes for the association of rs11591147 across the phenome (Figure 3). TreeWAS using a six-variant genetic score (Table 1) showed six active nodes, all of which were among the 11 active in the single-variant test (Figure S4).

**Figure 3:**
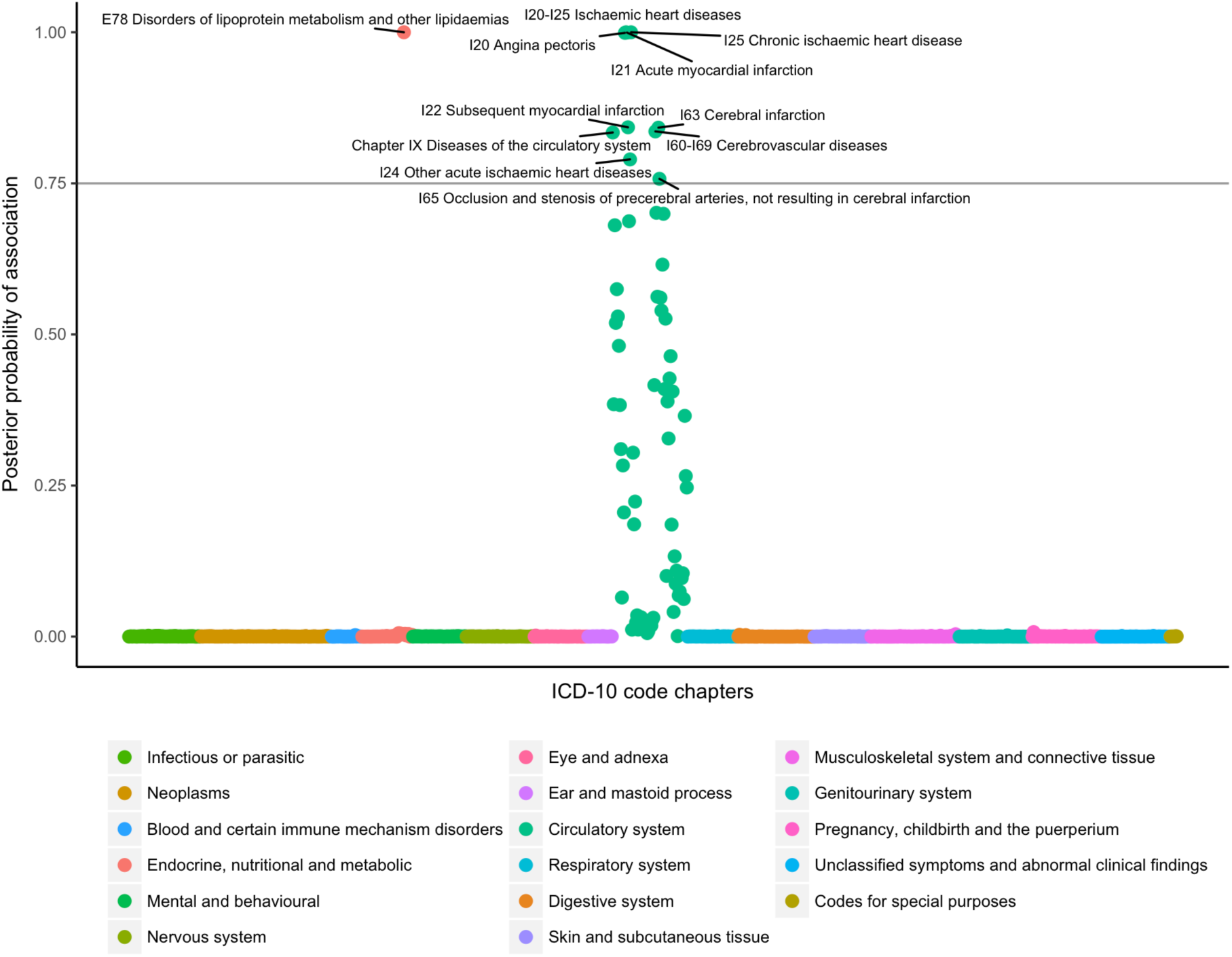
TreeWAS association results for rs11591147 (T) across the phenome. Associations with a posterior probability > 0.75 are considered significant.

## Discussion

In our phenome-wide analysis of 337,536 participants of the United Kingdom Biobank, we studied associations of LoF variants in *PCSK9* with 11-predefined phenotypes (based on prior literature) as well as across the whole phenome to discover potential opportunities for drug repositioning or risks of adverse drug effects. We made several important observations.

First, we observed robust and replicable associations of the *PCSK9* missense variant rs11591147 with cholesterol medication, coronary heart disease and ischemic stroke; the last has not been observed in prior large genetic studies.

Second, we did not observe associations of this variant with T2D, cataracts, epilepsy and cognitive dysfunction, all potential side effects of *PCSK9* inhibitors that have been suggested in prior literature; we did not see evidence suggesting these associations even after re-analyzing the full data set (statistical power in Table S1).

Third, using two alternative but related methods to study associations of *PCSK9* LoF variants across the phenome, we did not observe associations with phenotypes other than those associated with LDL-C and atherosclerosis, indicating that if there are additional side effects of *PCSK9* inhibition, they are likely to be of small effect sizes and/or rare. We observed nominal associations with 19 variables across the phenome, which were not significant using our *a priori* defined thresholds (Figure S3). *PCSK9* inhibitors can also have other monoclonal antibody related side effects that are difficult to test using this approach.

Phenome-wide association studies of gene-disrupting variants that influence the activity of drug target genes represent a promising strategy to predict drug effects, which can include adverse effects or potential drug repositioning opportunities. Such studies depend on the availability of relevant coding variants that can mimic pharmaceutical interventions; other variants associated with gene activity, such as expression quantitative trait loci (eQTLs), can also be used as proxies. Although randomized controlled trials are the gold standard for evidence of primary drug effects, they are very costly and time-consuming; genetic studies can provide preliminary evidence predicting the outcome of such studies. Further, side effects reported in clinical trials are sometimes equally likely in the placebo and treatment arms, can be influenced by factors other than primary drug action,^31^ and are often too rare to discover in Phase 3 trials. Studying gene disrupting variant effects in large genomic cohorts is therefore a complementary method to predict new indications and side effects associated with primary drug action. The development and clinical success of *PCSK9* inhibitors is one of the best examples of the value of genetics in drug discovery.^10,21,23,32^ Like studies on statin effects, several clinical trials and genetic studies have explored the effects of *PCSK9* inhibitors on lipid levels, coronary heart disease, mortality, stroke, T2D and cognitive effects; these studies are often hampered by small sample sizes and offer conflicting evidence that can impact our understanding of new indications, safety considerations, and side effects.^12,33–35^ Hence, we developed a list of 11 hypothesis-driven phenotypes with careful definitions to evaluate the evidence of association of *PCSK9* variants with phenotypes of interest in concurrent literature.^21,24,26,27^

Consistent with many prior studies,^14,17,21^ we observed a protective effect of the *PCSK9* LoF variant rs11591147 on cholesterol medication and coronary heart disease. Since lipid measurements have not been completed in the UKB, self-reported cholesterol medication served as a proxy response variable for high lipid levels. Using measured lipid levels is likely to increase power, so our effect estimate is likely to be biased towards the null. Effects of *PCSK9* inhibition (genetic and pharmacologic) on LDL-C levels and coronary heart disease are well-established.^25^ A post-hoc analysis of the treatment and control groups from the ODYSSEY LONG TERM trial showed a lower rate of adverse cardiovascular events in the treatment group.^36^ The cardiovascular impacts of *PCSK9* inhibitors was more formally addressed in the FOURIER^32^ trial, which confirmed the protective effects of the *PCSK9* inhibitor evolocumab on composite end points comprised of many cardiovascular outcomes. Genetic studies have also confirmed this protective effect on varied individual definitions and composite endpoints of adverse cardiovascular outcomes.^37^

There are conflicting results about the effects of *PCSK9* inhibitors on stroke, and only exploratory clinical evidence of this association is available. Given the large sample size in the UKB, we observed a significant protective association between rsII59H47 and ischemic stroke (OR = 0.61, p = 0.002) that replicated using our thresholds. We defined ischemic stroke to include infarction and occlusion of relevant blood vessels. In addition, we observed a nominally significant association with all stroke in the full dataset (OR = 0.82, p = 0.01), which was not significant in the discovery subset using our predefined criteria. We did not observe an association with hemorrhagic stroke (p = 0.81). TreeWAS single variant analysis showed associations with cerebral infarction and occlusion. Our results suggest that *PCSK9* inhibitors are likely to be protective against ischemic stroke, which is consistent with our knowledge about associations of LDL-C, atherosclerosis and ischemic stroke. Prior clinical studies, such as the ODYSSEY LONG TERM and FOURIER trials, have included ischemic stroke within more comprehensive endpoints or only addressed it in exploratory analyses. Non-fatal is-chemic stroke is included in the primary endpoint of the ongoing ODYSSEY OUTCOMES Phase 3 clinical trial. A handful of small studies have shown suggestive evidence for the association between *PCSK9* variants and ischemic stroke.^12,35,38^ However, larger studies have offered conflicting evidence. Two recent meta-analyses using genetic data^39^ and randomized clinical data^40^ showed no association between *PCSK9* inhibition and ischemic stroke. The association between clinical conditions resulting in altered lipid levels, such as FH, and stroke is unclear; prospective studies have shown no relationship between FH and stroke mortality, but show a relationship between FH and peripheral artery disease.^41^ One study showed that non-statin lipid-lowering interventions are not associated with a reduction in fatal stroke, but an update to the study with data from the Improved Reduction of Outcomes Vytorin Efficacy International Trial (IMPROVE-IT) suggested that all cholesterol-lowering interventions should reduce the risk of stroke.^42^ Our results using a Mendelian Randomization approach in the UKB support this hypothesis. The genetic study of PCSK9 variants by Hopewell et al.,^39^ which showed no association between rs11591147 and ischemic stroke in data from the METASTROKE Consortium, was a meta-analysis of 12 case-control studies with 10,307 cases and 19,326 controls; our study uses the UKB, a cohort study with 2,622 cases and 334,914 controls. Ischemic stroke was defined similarly using clinical diagnoses in both studies, but cases in some studies participating in METASTROKE were also confirmed using brain imaging. The difference in the observed effect on ischemic stroke in the two studies could be due to these slight differences in case definitions, variability among individual datasets included in the meta-analysis, difference in statistical power, or chance.

We did not observe an association of rs11591147 with T2D. Several studies have suggested that LDL-C concentrations have a causal link to an increase in T2D risk independent of lipid-lowering medication.^26,43^ Meta-analyses of unintended effects of statin therapy have shown an increase in T2D risk,^31^ and genetic studies of variants in *HMGCR* and *NPC1L1* (encoding the molecular targets of statins and ezetimibe, respectively) have shown increased risk of T2D associated with LDL cholesterol-lowering variants.^44^ A study of *PCSK9* variants showed a 6.1% (OR 1.06) increase in T2D risk in individuals with alleles protective against high LDL-C levels; the detected risk increased to 12.7% (OR 1.11) after adjustment for LDL-C levels.^34^ Another Mendelian Randomization study showed a similar increase in T2D risk among carriers of *PCSK9* variants.^25^ However, there is also conflicting evidence of this association. A pooled study of ten ODYSSEY Phase 3 trials showed no increase in new-onset diabetes between alirocumab treatment and control groups during a follow-up period of 6-18 months.^33^ A further analysis of the FOURIER evolocumab trial data showed that treatment resulted in a similar reduction in LDL-C levels in patients with and without prevalent diabetes, and did not worsen glycemia or increase the risk of new-onset diabetes.^21^ However, given the evidence of association between LDL-C reduction and T2D risk across treatment modalities, the lack of association between rs11591147 and T2D in the UKB could be due to the small effect size of *PCSK9* variants on T2D and lack of precision of the diabetes diagnosis in our study; both decrease power to detect a weak association. Indeed, prior genetic studies of *PCSK9* variants have including a larger number of T2D cases; 51,623 as compared to 20,947 in our study. Our power to detect an association with T2D was 52% for an OR of 1.11.^25^ Cataracts are known to be associated with T2D,^45^ and we did not observe an association between *PCSK9* inhibition and cataracts.

Prior studies have indicated that lowering LDL-cholesterol can alter the risk of epilepsy^46^ and cognitive dysfunction,^27^ but the evidence is very weak and inconsistent. Similar concerns have been raised for *PCSK9* inhibitors; the ongoing EBBINGHAUS trial, a cognitive study of patients enrolled in the FOURIER trial, is seeking to evaluate whether the addition of evolocumab to statin therapy impacts incident changes in cognition.^27^ We found no association between rs1159H47 and epilepsy or cognitive function measured using a trail-making test. We also did not observe significant associations among cognitive function variables included in the phenome-wide analyses, including dementia and Parkinson's disease diagnoses, but observed a nominal association with mood affective disorders (Figure S3).

*PCSK9* inhibitors comprise a promising new category of lipid-lowering therapy for CVD. We provide genetic evidence that, in addition to coronary heart disease, *PCSK9* inhibitors are likely to be protective against ischemic stroke, defined here using infarction and occlusion of relevant blood vessels. We did not find evidence of cognitive side effects of *PCSK9* inhibitors in the UKB; this result adds genetic support for the potential outcomes of the ongoing EBBINGHAUS trial. Finally, we also did not observe any associations with T2D; we were underpowered to detect a small effect of *PCSK9* variants on T2D as defined in UKB.

## Acknowledgements

The authors thank Stefan Gustafsson for assistance with data management. This research was conducted using the UK Biobank Resource and supported by a National Heart, Lung and Blood Institute (NHLBI) ROI HLI353I3-0I to E.I.

## References

1. Collaborators GBDCoD. Global, regional, and national age-sex specific mortality for 264 causes of death, 1980–2016: a systematic analysis for the Global Burden of Disease Study 2016. Lancet. 2017;390:1151–1210.

2. Mahmood SS, Levy D, Vasan RS and Wang TJ. The Framingham Heart Study and the epidemiology of cardiovascular disease: a historical perspective. Lancet. 2014;383:999–1008.

3. Nelson RH. Hyperlipidemia as a risk factor for cardiovascular disease. Prim Care. 2013;40:195–211.

4. Baigent C, Keech A, Kearney PM, Blackwell L, Buck G, Pollicino C, Kirby A, Sourjina T, Peto R, Collins R, Simes R and Cholesterol Treatment Trialists C. Efficacy and safety of cholesterol-lowering treatment: prospective meta-analysis of data from 90,056 participants in 14 randomised trials of statins. Lancet. 2005;366:1267–78.

5. Stone NJ, Robinson JG, Lichtenstein AH, Goff DC Jr., Lloyd-Jones DM, Smith SC Jr., Blum C, Schwartz JS and Panel AACG. Treatment of blood cholesterol to reduce atherosclerotic cardiovascular disease risk in adults: synopsis of the 2013 American College of Cardiology/American Heart Association cholesterol guideline. Ann Intern Med. 2014;160:339–43.

6. Expert Dyslipidemia P and Grundy SM. An International Atherosclerosis Society Position Paper: global recommendations for the management of dyslipidemia. J Clinb Lipidol. 2013;7:561–5.

7. Cholesterol Treatment Trialists C, Baigent C, Blackwell L, Emberson J, Holland LE, Reith C, Bhala N, Peto R, Barnes EH, Keech A, Simes J and Collins R.Efficacy and safety of more intensive lowering of LDL cholesterol: a meta-analysis of data from 170,000 participants in 26 randomised trials. Lancet. 2010;376:1670–81.

8. Santos RD, Waters DD, Tarasenko L, Messig M, Jukema JW, Chiang CW, Ferrieres J and Foody JM. A comparison of non-HDL and LDL cholesterol goal attainment in a large, multinational patient population: the Lipid Treatment Assessment Project 2. Atherosclerosis. 2012;224:150–3.

9. Robinson JG and Stone NJ.The 2013 ACC/AHA guideline on the treatment of blood cholesterol to reduce atherosclerotic cardiovascular disease risk: a new paradigm supported by more evidence. Eur Heart J. 2015;36:2110–2118.

10. Abifadel M, Varret M, Rabes JP, et al. Mutations in *PCSK9* cause autosomal dominant hypercholesterolemia. Nat Genet. 2003;34:154–6.

11. Cohen J, Pertsemlidis A, Kotowski IK, Graham R, Garcia CK and Hobbs HH.Low LDL cholesterol in individuals of African descent resulting from frequent nonsense mutations in *PCSK9*. Nat Genet. 2005;37:161–5.

12. Slimani A, Harira Y, Trabelsi I, Jomaa W, Maatouk F, Hamda KB and Slimane MN. Effect of E670G Polymorphism in *PCSK9* Gene on the Risk and Severity of Coronary Heart Disease and Ischemic Stroke in a Tunisian Cohort. J Mol Neurosci. 2014;53:150–7.

13. Timms KM, Wagner S, Samuels ME, Forbey K, Goldfine H, Jammulapati S, Skolnick MH, Hopkins PN, Hunt SC and Shattuck DM. A mutation in *PCSK9* causing autosomal-dominant hypercholesterolemia in a Utah pedigree. Hum Genet. 2004;114:349–53.

14. Cohen JC, Boerwinkle E, Mosley TH Jr. and Hobbs HH. Sequence variations in *PCSK9*, low LDL, and protection against coronary heart disease. N Engl J Med. 2006;354:1264–72.

15. Sanna S, Li B, Mulas A, et al. Fine mapping of five loci associated with low-density lipoprotein cholesterol detects variants that double the explained heritability. PLoS Genet. 2011;7:e1002198.

16. Wu Y, Waite LL, Jackson AU, et al. Trans-ethnic fine-mapping of lipid loci identifies population-specific signals and allelic heterogeneity that increases the trait variance explained. PLoS Genet. 2013;9:e1003379.

17. Surakka I, Horikoshi M, Magi R, et al. The impact of low-frequency and rare variants on lipid levels. Nat Genet. 2015;47:589–97.

18. Rashid S, Curtis DE, Garuti R, Anderson NN, Bashmakov Y, Ho YK, Hammer RE, Moon YA and Horton JD.Decreased plasma cholesterol and hypersensitivity to statins in mice lacking *PCSK9*. Proc Natl Acad Sci USA. 2005;102:5374–9.

19. Feng Q, Wei WQ, Chung CP, Levinson RT, Bastarache L, Denny JC and Stein CM. The effect of genetic variation in *PCSK9* on the LDL-cholesterol response to statin therapy. Pharmacogenomics J. 2017;17:204–208.

20. Abifadel M, Elbitar S, El Khoury P, Ghaleb Y, Chemaly M, Moussalli ML, Rabes JP, Varret M and Boileau C.Living the *PCSK9* adventure: from the identification of a new gene in familial hypercholesterolemia towards a potential new class of anticholesterol drugs. Curr Atheroscler Rep. 2014;16:439.

21. Sabatine MS, Giugliano RP, Keech AC, Honarpour N, Wiviott SD, Murphy SA, Kuder JF, Wang H, Liu T, Wasserman SM, Sever PS, Pedersen TR, Committee FS and Investigators. Evolocumab and Clinical Outcomes in Patients with Cardiovascular Disease. N Engl J Med. 2017;376:1713–1722.

22. Cannon CP, Cariou B, Blom D, McKenney JM, Lorenzato C, Pordy R, Chaudhari U, Colhoun HM and Investigators OCI. Efficacy and safety of alirocumab in high cardiovascular risk patients with inadequately controlled hypercholesterolaemia on maximally tolerated doses of statins: the ODYSSEY COMBO II randomized controlled trial. Eur Heart J. 2015;36:1186–94.

23. Giugliano RP and Sabatine MS. Are *PCSK9* Inhibitors the Next Breakthrough in the Cardiovascular Field? J Am Coll Cardiol. 2015;65:2638–51.

24. Sullivan D, Olsson AG, Scott R, Kim JB, Xue A, Gebski V, Wasserman SM and Stein EA. Effect of a monoclonal antibody to *PCSK9* on low-density lipoprotein cholesterol levels in statin-intolerant patients: the GAUSS randomized trial. JAMA. 2012;308:2497–506.

25. Schmidt AF, Swerdlow DI, Holmes MV, et al. *PCSK9* genetic variants and risk of type 2 diabetes: a mendelian randomisation study. Lancet Diabetes Endocrinol. 2017;5:97–105.

26. Ingelsson E and Knowles JW. Leveraging Human Genetics to Understand the Relation of LDL Cholesterol with Type 2 Diabetes. Clin Chem. 2017;63:1187–1189.

27. Giugliano RP, Mach F, Zavitz K, Kurtz C, Schneider J, Wang H, Keech A, Pedersen TR, Sabatine MS, Sever PS, Honarpour N, Wasserman SM, Ott BR and Investigators E. Design and rationale of the EBBINGHAUS trial: A phase 3, double-blind, placebo-controlled, multicenter study to assess the effect of evolocumab on cognitive function in patients with clinically evident cardiovascular disease and receiving statin background lipid-lowering therapy-A cognitive study of patients enrolled in the FOURIER trial. Clin Cardiol. 2017;40:59–65.

28. Demidenko E. Sample size determination for logistic regression revisited. Stat Med. 2007;26:3385–97.

29. Cortes A, Dendrou CA, Motyer A, Jostins L, Vukcevic D, Dilthey A, Donnelly P, Leslie S, Fugger L and McVean G. Bayesian analysis of genetic association across tree-structured routine healthcare data in the UK Biobank. Nat Genet. 2017;49:1311–1318.

30. Wu MC, Lee S, Cai T, Li Y, Boehnke M and Lin X. Rare-variant association testing for sequencing data with the sequence kernel association test. Am J Hum Genet. 2011;89:82–93.

31. Finegold JA and Francis DP. What proportion of symptomatic side-effects in patients taking statins are genuinely caused by the drug? A response to letters. Eur J Prev Cardiol. 2015;22:1328–30.

32. Sabatine MS, Giugliano RP, Wiviott SD, Raal FJ, Blom DJ, Robinson J, Ballantyne CM, Somaratne R, Legg J, Wasserman SM, Scott R, Koren MJ, Stein EA and Open-Label Study of Long-Term Evaluation against LDLCI. Efficacy and safety of evolocumab in reducing lipids and cardiovascular events. N Engl J Med. 2015;372:1500–9.

33. Colhoun HM, Ginsberg HN, Robinson JG, Leiter LA, Muller-Wieland D, Henry RR, Cariou B, Baccara-Dinet MT, Pordy R, Merlet L and Eckel RH. No effect of *PCSK9* inhibitor alirocumab on the incidence of diabetes in a pooled analysis from 10 ODYSSEY Phase 3 studies. Eur Heart J. 2016;37:2981–2989.

34. Ference BA, Robinson JG, Brook RD, Catapano AL, Chapman MJ, Neff DR, Voros S, Giugliano RP, Davey Smith G, Fazio S and Sabatine MS. Variation in *PCSK9* and *HMGCR* and Risk of Cardiovascular Disease and Diabetes. N Engl J Med. 2016;375:2144–2153.

35. Au A, Griffiths LR, Cheng KK, Wee Kooi C, Irene L and Keat Wei L. The Influence of *OLR1* and *PCSK9* Gene Polymorphisms on Ischemic Stroke: Evidence from a Meta-Analysis. Sci Rep. 2015;5:18224.

36. Hassan M. OSLER and ODYSSEY LONG TERM: *PCSK9* inhibitors on the right track of reducing cardiovascular events. Glob Cardiol Sci Pract. 2015;2015:20.

37. Scartezini M, Hubbart C, Whittall RA, Cooper JA, Neil AH and Humphries SE. The *PCSK9* gene R46L variant is associated with lower plasma lipid levels and cardiovascular risk in healthy U.K. men. Clin Sci (Lond). 2007;113:435–41.

38. Abboud S, Karhunen PJ, Lutjohann D, Goebeler S, Luoto T, Friedrichs S, Lehtimaki T, Pandolfo M and Laaksonen R. Proprotein convertase subtilisin/kexin type 9 (*PCSK9*) gene is a risk factor of large-vessel atherosclerosis stroke. PLoS One. 2007;2:e1043.

39. Hopewell JC, Malik R, Valdes-Marquez E, Worrall BB, Collins R and ISGC MCot. Differential effects of *PCSK9* variants on risk of coronary disease and ischaemic stroke. Eur Heart J. 2017.

40. Milionis H, Barkas F, Ntaios G, Papavasileiou V, Vemmos K, Michel P and Elisaf M. Proprotein convertase subtilisin kexin 9 (*PCSK9*) inhibitors to treat hypercholesterolemia: Effect on stroke risk. Eur J Intern Med. 2016;34:54–57.

41. Hutter CM, Austin MA and Humphries SE. Familial hypercholesterolemia, peripheral arterial disease, and stroke: a HuGE minireview. Am J Epidemiol. 2004;160:430–5.

42. Castilla-Guerra L and Fernandez-Moreno MC. *PCSK9* inhibitors: A new era in stroke prevention? Eur J Intern Med. 2017;37:e44.

43. Fall T, Xie W, Poon W, Yaghootkar H, Magi R, Consortium G, Knowles JW, Lyssenko V, Weedon M, Frayling TM and Ingelsson E. Using Genetic Variants to Assess the Relationship Between Circulating Lipids and Type 2 Diabetes. Diabetes. 2015;64:2676–84.

44. Lotta LA, Sharp SJ, Burgess S, et al. Association Between Low-Density Lipoprotein Cholesterol-Lowering Genetic Variants and Risk of Type 2 Diabetes: A Meta-analysis. JAMA. 2016;316:1383–1391.

45. Klein BE, Klein R and Moss SE. Prevalence of cataracts in a population-based study of persons with diabetes mellitus. Ophthalmology. 1985;92:1191–6.

46. Benn M, Nordestgaard BG, Frikke-Schmidt R and Tybjaerg-Hansen A. Low LDL cholesterol, *PCSK9* and *HMGCR* genetic variation, and risk of Alzheimer’s disease and Parkinson’s disease: Mendelian randomisation study. BMJ. 2017;357:j1648.

